# Prey attracting but not avoiding predators suggests an asymmetric investment in the predation sequence

**DOI:** 10.1101/2024.11.13.623348

**Authors:** Nicolas Ferry, Christian Fiderer, Anne Peters, Axel Ballmann, Marco Heurich

## Abstract

Understanding predator-prey interactions, particularly how species use space and time to influence encounter rates, is fundamental in community and behavioural ecology. However, for large, free-ranging animals, encounter rates are rarely quantified directly, because of logistical and methodological challenges associated with tracking both predators and prey simultaneously. Instead, studies commonly rely on proxies such as spatial or temporal overlap. While informative, these proxies provide only incomplete estimates of encounter rates because they typically consider only one dimension of the interaction process (spatial or temporal). Although camera traps cannot directly measure encounters among large free-roaming species, they offer the opportunity to quantify the proximal co-occurrence, i.e., the extent to which species tolerate or avoid one another’s proximity in space and time. We analysed data from a one-year study conducted across four German protected areas using 283 camera traps and applying recurrent event analysis to investigate interactions among three prey species, red deer (*Cervus elaphus)*, roe deer (*Capreolus capreolus)*, wild boar (*Sus scrofa*) and two large predators grey wolf (*Canis lupus)*, and Eurasian lynx (*Lynx lynx*). Prey visitation rates were unaffected by predator presence, whereas wolves exhibited a strong attraction to prey, with visitation rates four to six times higher immediately after prey detections. The limited sample size prevented robust conclusions regarding Eurasian lynx responses. These findings point towards an asymmetry in the predation sequence (i.e., spatio-temporal proximity, encounter, ignorance or avoidance post-encounter, capture or escape from attack): Predators must succeed at every stage of the sequence to capture prey, while prey can avoid predation by disrupting the process at any single stage. Our results suggest that prey species do not necessarily reduce large-scale spatio-temporal proximity to predators and may instead rely on anti-predator responses occurring later in the predation sequence.

## Introduction

Understanding predator-prey interactions is fundamental to ecology, with implications for individual fitness, population dynamics, community structure, and ecosystem functions (Sih et al. 1985; Krebs et al. 1995). Holling (1959) as well as Lima & Dill (1990) conceptualized predation as a sequence of successive steps leading to the predatory event: (a) spatio-temporal proximity, (b) encounter, (c1) ignorance or (c2) avoidance post-encounter or eventually (c3) the attack leading either to (d1) the escape or (d2) the capture of the prey, i.e. the predation event. A key component is therefore the rate of encounters, i.e., when the detection range of the predator or prey exceeds the distance between them (*sensu* Lima & Dill, 1990). All other factors being equal, prey should seek to maximize use of safer areas and periods, following a dynamic landscape of fear (Messier & Barrette, 1985; Laundré et al. 2001, Palmer et al. 2017), i.e., the spatio-temporal variation of predation risk perceived by prey (Gaynor et al. 2019), in order to minimize the encounter rate. On the other hand, optimal foraging predicts that predators, seeking to maximize the encounter rate should forage where there is the highest chance of finding and capturing prey (Sih, 1980). These two opposite interests, i.e. the “space race” (Sih, 1984), can lead to counter-intuitive patterns except when considering both predator and prey simultaneously (e.g. Laundré et al. 2009, Kuijper et al. 2013). Further, the physical landscape affects predator and prey spatial and temporal use, in part due to energy expenditure (Dickinson et al., 2000) but also due to their perceived probability of encountering predators/prey. Prey may choose habitats reducing encounter rates with predators or lowering death probabilities upon encounters (Heithaus et al., 2009). However, how the environment shapes the predation interaction is complex. For instance, vegetation density can either provide safety or act as an obstacle, depending on context and species (Hernandez & Laundré, 2005; Ordiz et al., 2011; Camp et al., 2013, Kuijper et al. 2013).

Estimating this encounter rate is challenging, especially for large free-ranging animals (Hebblewhite et al., 2005, Hoffman et al., 2022). Traditionally, spatial overlap has been the primary proxy used (e.g., occupancy, MacKenzie et al., 2004; resource selection probability, Hebblewhite et al., 2005; habitat use similarity, Smith et al., 2019a). However, space overlap alone is insufficient to approximate the encounter rate (Suraci et al. 2022), which depends on predator and prey biology (e.g., population density, foraging tactic, locomotion, sensory abilities) as well as environmental conditions (Sih, 1984; Lima & Dill, 1990). Even with complete spatial overlap, low population densities (Travis & Palmer, 2005; Sims et al., 2006) or slower movement speeds (Sih, 1984) can reduce encounter rates. Additionally, predators and prey must co-occur in both space and time for encounters to happen (Sih, 1984; Minta, 1992; Caro, 2005; Sih, 2005; Suraci et al., 2022). Temporal overlap within cycles (e.g., circadian) can serve as another proxy for encounter rates, facilitated by analytical advancements (e.g., diel activity pattern overlap, Ridout & Linkie, 2009) and technologies (e.g., GPS collars, Eriksen et al., 2011; camera traps, Harmsen et al., 2009). However, summarizing linear time in a circular pattern (yearly or daily) only provides a partial view. High temporal and spatial overlap may occur, but individuals can avoid encounters by consistently being out of sync by a day or more. Further, interpreting these overlaps can be challenging with regard to the race between prey and predator without proper experimental design (e.g., in presence/absence of the predator). Thus, considering spatial and temporal overlaps separately is insufficient for understanding how predators and prey aim to win the predation game. With large free-roaming species, telemetry is certainly the most accurate method for estimating encounter rates and attraction/avoidance processes (e.g., Li et al., 2013, Latombe et al., 2014). However, there are few empirical studies on this topic (e.g., Courbin et al., 2013; Middleton et al., 2013; Isbell et al., 2018; Tallian et al., 2023; Elbroch et al., 2017; Rafiq et al., 2020). This scarcity is likely due to the difficulty of simultaneously monitoring two large free-roaming species with sufficient sample sizes and defining an adequate detection radius to identify encounters (Lynch et al., 2015, Latombe et al., 2014). Stationary monitoring devices, like camera traps, cannot accurately estimate encounter rates because they rarely capture direct encounters (Triguero-Ocaña et al., 2020, Suraci et al., 2022), except in specific situations such as predation on corralled livestock (e.g., Hoffman et al., 2022). However, they can assess proximate co-occurrence, or the likelihood of one species being detected after another at a location within a certain time frame, indicating whether species tolerate or avoid each other (Schliep et al., 2018). Thus, camera traps can help bridge the gap between spatial/temporal overlaps and encounter rates (Suraci et al., 2022).

Grey wolf (*Canis lupus*) and Eurasian lynx (*Lynx lynx*) populations are recovering in western and central Europe after near-extinction (Chapron et al., 2014). Wolves are cursorial predators, relying on long-distance movements (Mech, 1970; Peterson & Ciucci, 2003). Red deer (*Cervus elaphus*), roe deer (*Capreolus capreolus*) and wild boar (*Sus scrofa*) are a significant part of the wolves’ diet (Newsome et al., 2016). Lynx are ambush predators, mainly preying on roe deer in central Europe, occasionally on red deer and rarely on wild boar (Khorozyan & Heurich, 2023; Okarma et al., 1997; Molinari-Jobin et al., 2007; Krofel et al., 2011; Heurich et al., 2016). Because of the recolonisation of European landscapes by wolf and lynx, these three herbivores face an increasing risk of non-human predation, providing new opportunities to examine predator-prey interactions in European ecosystems (see Gerber et al. 2024 for a review on wolf non-consumptive effects). From a prey perspective, apart from human-induced mortality, wolf predation is the main cause of death for red deer while lynx predation is less significant, with lynx preying mainly on juvenile and female red deer (Kamler et al., 2007; Okarma et al., 1995; Belotti et al., 2015). Roe deer is similarly predated by both lynxes and wolves, whereas wild boar is primarily predated by wolves and almost never by lynx (Okarma et al., 1995). Using camera trap data from natural systems in which the three ungulate species and at least one of the large predators occur represents therefore an ideal situation to assess how animals are using space and time according to different levels of predation pressure.

In this study, we used camera traps to investigate the proximate co-occurrence between the two predators (wolf, lynx) and the three herbivores (red deer, roe deer and wild boar). Based on prey prevalence in a predator’s diet, predator impact on prey mortality, and prey responses to predator hunting strategies (cursorial vs. ambush, Kuijper et al., 2014; Sunde et al., 2022), we hypothesized various scenarios as follows (see Fig. 1 for a visual depiction of the hypotheses):

(H1). Red deer show avoidance towards wolves and, to a lesser extent, towards lynx.
(H2). Roe deer show avoidance towards both wolves and lynx.
(H3). Wild boar show avoidance towards wolves and remain neutral towards lynx.
(H4). Wolves are attracted to all three herbivores.
(H5). Lynx are attracted to roe deer, less attracted to red deer, and neutral towards wild boars. Further, information on previous species presence is mandatory for observing a reactive avoidance or attraction effect. Olfactory clues like fur, skin, feces, or urine, are recognized as important signals for heterospecific local presence (Kuijper et al., 2014; Parsons et al., 2018, Lawson et al. 2019 and references therein) and are assumed to diminish over a few days or a week (Lima, 1998; Parsons et al., 2018; Prat-Guitart et al., 2020). Consequently, we anticipate that all visitation rates will return to baseline levels after a few days (Fig. 1). Finally, considering the influence of the landscape on predator-prey interactions, we hypothesize that:
(H6). Prey avoidance will be higher towards wolves in open areas and towards lynx in denser areas.
(H7). Predator attraction will be higher towards prey in open areas for wolves and in denser areas for lynx.

**Figure 1:**
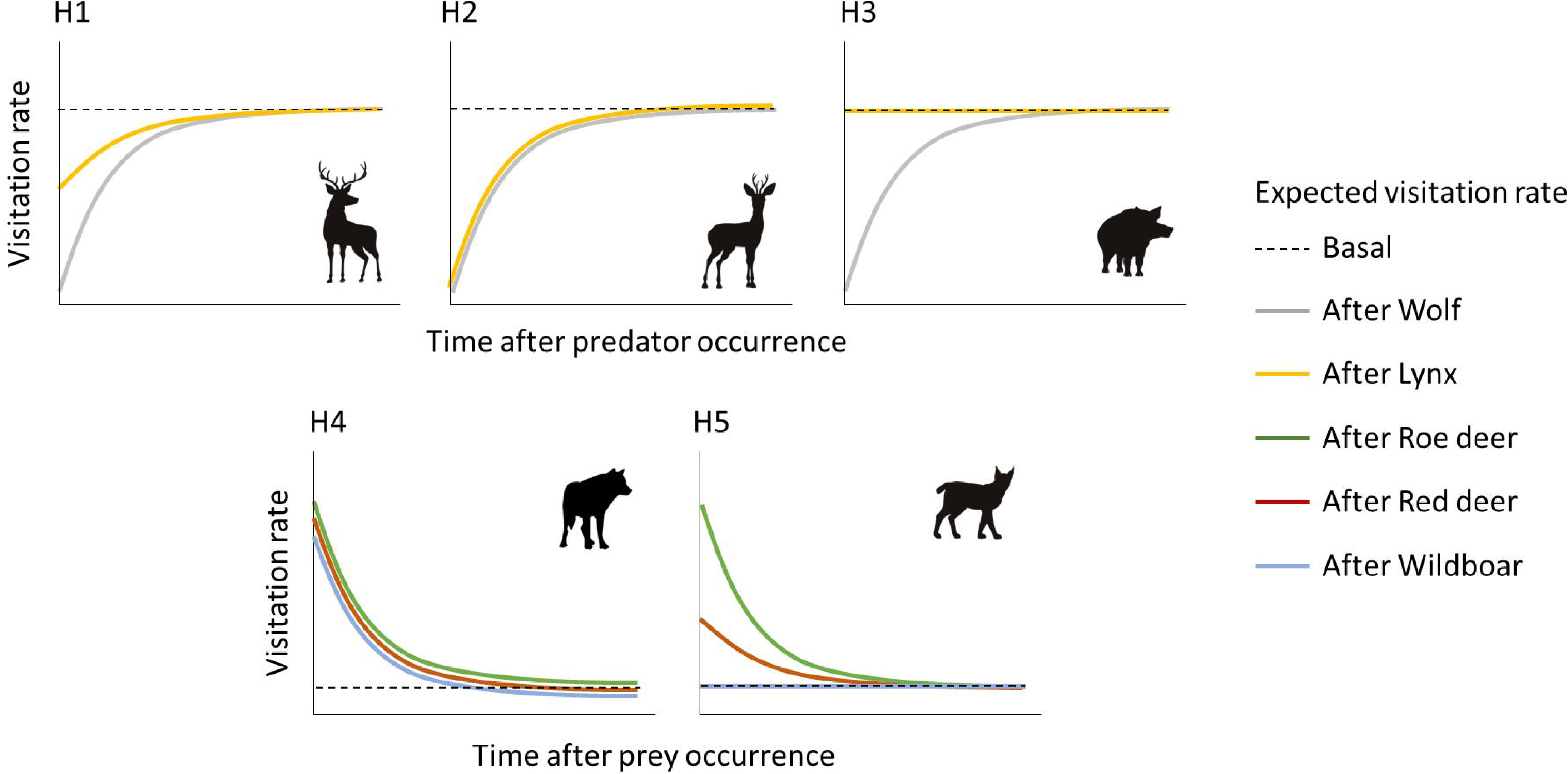
diagram of the expected visitation rate dynamics over time based on hypotheses H1-H5. The pattern depends on the effects type (avoidance, neutral, attraction) and is relative to the baseline visitation rate, i.e., under “usual” circumstances: 1) avoidance, starting lower than baseline level and increasing until coming back to this last (H1-H3), 2) neutral, no change from the baseline level (H3 and H5) and 3) attraction, starting higher than baseline level and decreasing until coming back to this latter (H4-H5).

## Methods

### Study sites

Camera trap data was collected from four large, protected areas in Germany: Bavarian Forest National Park (BFNP), Müritz National Park (MNP), Harz National Park (HNP), and Königsbrücker Heide wilderness area (KHWA, Fig. 2). Due to the protection status of the study areas, human activities are restricted to recreation, hunting, and forestry operations necessary to meet area-specific management objectives. Across all areas, the primary goal of hunting is to reduce browsing pressure and minimize conflicts arising from browsing and crop damage outside the protected areas, thereby reducing conflicts with neighboring landowners, such as farmers and foresters. During the study period, in HNP, hunting of red deer, roe deer, and wild boar takes place from August to December throughout the entire park, using both drive hunts and individual hunting. In MNP, hunting is permitted in 35% of the area and is conducted through drive hunts and individual hunting. Red deer and roe deer are hunted from August to January, with male and yearling roe deer additionally hunted in May. The hunting season for fallow deer extends from September to January, while wild boar are hunted year-round. In the BFNP, hunting is limited to the management zone covering 25% of the park’s border area. Here, individual hunting and baiting are practiced. Red deer are hunted from June to January, and wild boar are hunted year-round. In the KHWA, single hunt and drive hunts are permitted in the bordering areas (ca. 19 percent of the total protected area). Hunting is mainly focused on red deer, fallow deer (both from August to January), roe deer (males and young adults from mid-April to January and females and fawns from August to January) and wild boar (year-round). These areas were sampled in the project “Ungulate Monitoring in German National Parks” from 2019 to 2020.

**Figure 2:**
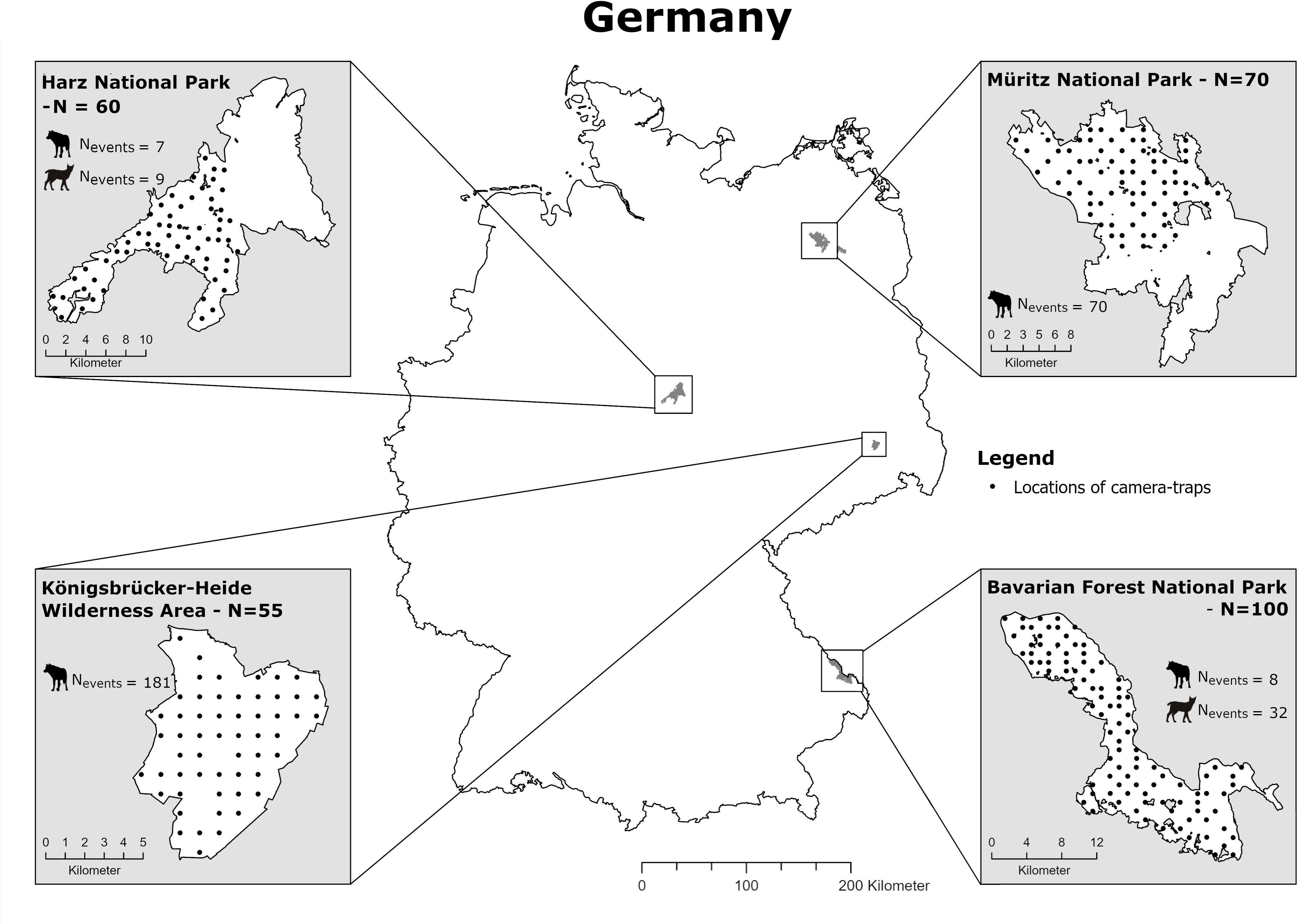
Map of the four protected areas monitored in Germany from October 2019 to October 2020. The number of camera traps deployed in each area is specified next to the area’s name (N). Black dots indicate camera trap locations. The number of independent wolf and lynx events is provided for each area (N_events_).

Lynx were reintroduced to the BFNP during the 1970s and 1980s (Wölfl et al., 2001). In the HNP, reintroduction took place between 2000 and 2006, with animals originating from German and Swedish zoos and wildlife parks (Mueller et al., 2020). Natural dispersal and the reestablishment of lynx in currently unoccupied areas across Germany are hindered by habitat fragmentation, large distances between populations, road mortality, and likely illegal killings outside protected areas (Kramer-Schadt et al., 2004; Magg et al., 2016). As no reintroduction efforts have been undertaken in MNP and KHWA, these areas are currently not inhabited by lynx. Wolves naturally recolonized Germany since the early 2000s, with wolves migrating from Eastern Europe, and were present in all areas during the time data was collected (Jarausch et al., 2021).

### Camera-trap deployment

A total of 285 camera traps were deployed between October 1, 2019, and October 1, 2020, resulting in 91,625 camera trapping days (Table 2). When calculating trapping days, periods during which cameras were inactive or unable to record images—e.g., due to snow covering the lens—were excluded. In all study areas we used Cuddeback C2 cameras equipped with an infra-red flash. In MNP, Reconyx Hyperfire cameras were used at ten additional and randomly selected locations, while in BFNP, we used 50 Bushnell Trophy Cam cameras in the northern area and 50 Cuddeback C2 in the southern area. In July, all 100 cameras in the BFNP were replaced with Cuddeback G models.

**Table 1:** study sites description. *m.a.s.l.: meter above sea level. **Definition of common mammal species was done based on the study camera p monitoring.

| Study area | Size (km<sup>2</sup>) | Hunting free area (%) | Elevation (m.a.s.l.*) | Overall habitat type | Main habitat | Common mammal species** |
| --- | --- | --- | --- | --- | --- | --- |
| **BFNP** | 243 | 75% | 600 - 1,453 | Low mountain range | Alluvial spruce-, mixed alpine-, mountain spruce forest (Kreft 2011) | Red deer, roe deer, wild boar, Eurasian lynx, red fox, badger, pine marten |
| **HNP** | 247 | 0 | 200 - 1,141 | Low mountain range | Mountain spruce-, mixed montane-, beech forests (Kreft 2011) | Red deer, roe deer, wild boar, Eurasian lynx, red fox, badger, |
| **MNP** | 322 | 65% | 62 - 143 | lowland | Sub-alpine dwarf shrubland/moors/running water and rock biotopes (Kreft 2011) | Red deer, roe deer, wild boar, fallow deer, wolf, red fox, badger, racoon, hare |
| **KHWA** | 69 | 81% | 100 - 160 | lowland | Riparian forest, pine bog forests, heaths, grassland, mires | Red deer, roe deer, wild boar, wolf, red fox, badger, hare |

**Table 2:** camera trap deployment information per study site. Range of CT days represents the minimum and maximum number of days a camera trap was active. CT days is the accumulated camera trap days.

| National Park | Cameras | Camera model | Range of CT days | CT days |
| --- | --- | --- | --- | --- |
| **BFNP** | 100 | 50 Cuddeback C2 +<br>50 Bushnell Trophy Cam /<br>100 Cuddeback G | 29 - 367 | 30,576 |
| **HNP** | 60 | Cuddeback C2 | 50 – 367 | 19,444 |
| **MNP** | 70 | Cuddeback C2 +<br>10 Reconyx | 7 - 367 | 22,377 |
| **KHWA** | 55 | Cuddeback C2 | 283 - 367 | 19,228 |

Camera trap locations were selected using a systematic random sampling approach based on a randomly placed grid with a cell size of 1×1 km, where the center of each grid cell served as a potential camera location. Cameras were therefore not deliberately placed along game trails or other features that might increase the likelihood of wildlife detection (Wearn & Glover-Kapfer, 2017). The 1×1 km grid cell size was chosen to ensure spatial independence among camera trap locations (Wearn & Glover-Kapfer, 2017). Grid cells were excluded prior to final selection if (1) their centers did not overlap with the study area, or (2) if the grid cell center was located in an area unsuitable for camera deployment, such as water bodies, steep slopes, or fenced areas. From the remaining grid cells, 50–100 were randomly and evenly selected across each study area using ArcGIS software. In cases where the number of available cameras within a national park was insufficient to cover the entire area - such as in HNP and MNP - the park was divided into two sub-areas, and only one sub-area was surveyed. Consequently, in HNP, cameras were deployed only in the section located within the federal state of Lower Saxony (western part), while in MNP, only the northern section was surveyed (Fig. 1). Camera traps were installed within 25 m of the designated grid cell center, mounted either on trees or on wooden posts in open habitats. The lens height was set at 50 cm to ensure effective detection of medium- to large-sized mammals. Cameras were positioned to provide at least 8 m of unobstructed view along the line of sight and oriented northward (±10°) to guarantee conscious orientation and minimize reflections from direct sunlight. All camera traps were configured to capture three consecutive still images with minimal delay upon activation. Video mode was not used.

### Vegetation density

In August 2019, vegetation density around each camera trap was measured. To assess vegetation density, a stake was placed 8 m in front of the camera along its visual axis. Four additional markers were placed at 4 m and 12□m along the camera’s visual axis, and 4□m west and east of the centre, forming a square around the central point with 90° angles. Vegetation density was assessed at the four outer markers. A red towel was stretched from each marker toward the centre and photographed at 100□cm lens height in landscape format, using automatic mode without zoom or flash. The towel was held taut and parallel to the ground, with edges aligned to the image frame; camera angles were adjusted on slopes as needed. Inaccessible markers were visually estimated, and if obstacles prevented towel placement, it was positioned at the nearest feasible location, with the adjusted distance recorded (Abrams et al., 2018). Pictures were cropped to the extent of the red blanket and converted to black and white using the image processing software Gimp2 (https://gimp.org). Vegetation density was calculated as the ratio of black to white pixels, averaged across all four photos (0 = completely open, 1 = completely closed). Analysis of the vegetation pictures took place with the R-package *imageSeq* (Niedballa et al., 2022).

### Camera trap data preparation and “survey” creation

All data preparations and analyses were done using R 4.3.0 open-source software (R Core Team, 2023). Each event was a detection of either a single individual or a cluster of individuals. Multiple detections of the same species and at the same location less than five minutes apart were considered as a single independent event (Henrich et al., 2022). Following Ferry et al. (2024) approach, when quantifying for instance the effect of wolf on red deer visitation rate over time, wolf and red deer were designated as the *primary* (the one affecting) and *secondary* (the one affected) species, respectively (*sensu* Niedballa et al. 2019, Ferry et al., 2024). Whenever a wolf, the *primary*, was observed at a site, a new “post-primary survey” (*survey* hereafter) commenced at that site and continued for a defined period (details on survey duration below). Throughout this survey period, all observations of red deer, the *secondary* species, were recorded and treated as *recurrent* events (Appendix 1 - Fig. S1). It was acknowledged that additional species might also influence the secondary visitation rates, either through attraction or avoidance. To mitigate potential confounding effects from these species when assessing the impact of *primary* on *secondary* visitation rates, we established a *censoring* species pool (Ferry et al. 2024). This censoring pool consisted of the primary species and *tertiary* species—those other than the primary assumed to affect the secondary species through avoidance or attraction. Therefore, if we assessed the effect of wolf on red deer visitation rate, wolf was the *primary*, red deer was the *secondary* and lynx and human were considered as *tertiary* species potentially affecting red deer visitation rate. The occurrence of any of these three censoring species (wolf, lynx, human) ended the current survey (Fig. S1 A). If we assessed the effect of wild boar (*primary*) on lynx (*secondary*), the censoring pool comprised wild boar but also red deer, roe deer and human (*tertiaries*). Camera traps sometimes failed due to issues such as empty batteries or snow obstruction. Extended inactivity periods increase the chance of missing species observations, leading to an underestimation of the event rate. Inactivity periods were defined as survey “cut-offs” based on i) deployment data indicating camera trap daily activity and ii) the mean duration (in days) between two independent animal observations at the camera trap site. If an inactivity period exceeded the average duration between two events, it was considered a cut-off due to a high chance of missing animal observations, concluding the survey (Appendix 1 - Fig. S1 B). A survey could run for up to 300 days. However, considering olfactory cues are assumed to diminish over time (Prat-Guitart et al., 2020; Lima, 1998; Parsons et al., 2018), we set an arbitrary maximum survey duration of 30 days. This timeframe balances capturing primary’s effect on the secondary visitation rate and ensuring model estimation is not solely influenced by unrelated factors (e.g., resource availability phenology). As a summary, the survey ended with one of the following censoring events: i) another species from the censoring pool observed (primary or tertiary), ii) prolonged camera trap inactivity (“cut-off” period), iii) the survey reached a 30-day duration threshold or iv) study’s end.

The transformation of raw camera trap data into the recurrent event format was done using the *recurrent* function from the *ct.recurrent* package (Ferry et al. 2024). For each recurrent event, the time since the start of each survey (t.stop) and the time of the previous recurrent event since the survey’s start (t.start, equals 0 if it was the first recurrent event) were determined. We recorded the event number and event status (0 = censoring event, 1 = recurrent event). A second data transformation step was required, from the recurrent event format, to obtain piece-wise exponential data using the *as_ped* function from the *pammtools* package (Bender et al. 2018). Data first have to be discretized into J intervals with cutpoints 0 = τ_0_ < τ_1_ < · · · < τ_J_. The *j*^th^ interval is defined as (τ_J−1_, τ_J_] and τ_J_ as the maximum time set arbitrarily, in this case, 30 days or less. Pseudo-observations were then created for each interval during which the survey was at risk of an event (Bender et al. 2018; Ramjith et al. 2021, see an example with ecological data in Ferry et al. 2024).

### Analysis

We then applied the Piece-wise Additive Mixed Model (PAMM, Bender & Scheipl, 2018; Ramjith et al. 2022) to estimate the temporal dynamics of the secondary species event rate after the passage of the primary species and tested an effect of the vegetation density on these dynamics. This approach was shown to be more efficient than other common methods in studying proximate co-occurrence (Ferry et al. 2024) such as the avoidance-attraction ratio (Niedballa et al. 2019, Dymit et al. 2025). As part of the GAM family, this model estimates linear and non-linear covariate effects using smoothing functions (see Ramjith et al., 2022 for a more detailed mathematical description). It enables the non-parametric estimation of the effect of time since the last primary event on the secondary’s visitation rate. Separate models were conducted for each prey-predator and predator-prey combination. We hypothesized different visitation rate dynamics according to vegetation density and tested this by examining the interaction between vegetation density and time after primary using bivariate penalized splines represented by tensor products (e.g. Ramjith et al. 2022, Ferry et al. 2024). We also incorporated the time of the day (*Sun Time*) and day of year (*DoY*) as control variables to account for circadian and seasonal dynamics of species event rates. Because animal activity is often attuned to sunrise and sunset, the timing of which varies with season, we used the average anchoring on the time of the day variable, from the activity package (Vazquez et al. 2019) to avoid the seasonal confounding effect. Due to sample size limitation, for few prey- predator combinations some of the variables/interactions were not included in the models (Table 3). All models included a survey’s random effect to address dependency between recurrent events of the same survey (*survey_id*), as well as a random effect of the National Park (*NP*) in order to encompass management difference, such as hunting, between them. The effect of smooth functions is expressed in estimated degrees of freedom (EDF), indicating the curvature’s complexity. The p-values test whether the smooth function significantly deviates from a flat line (Wood, 2017).

**Table 3:**
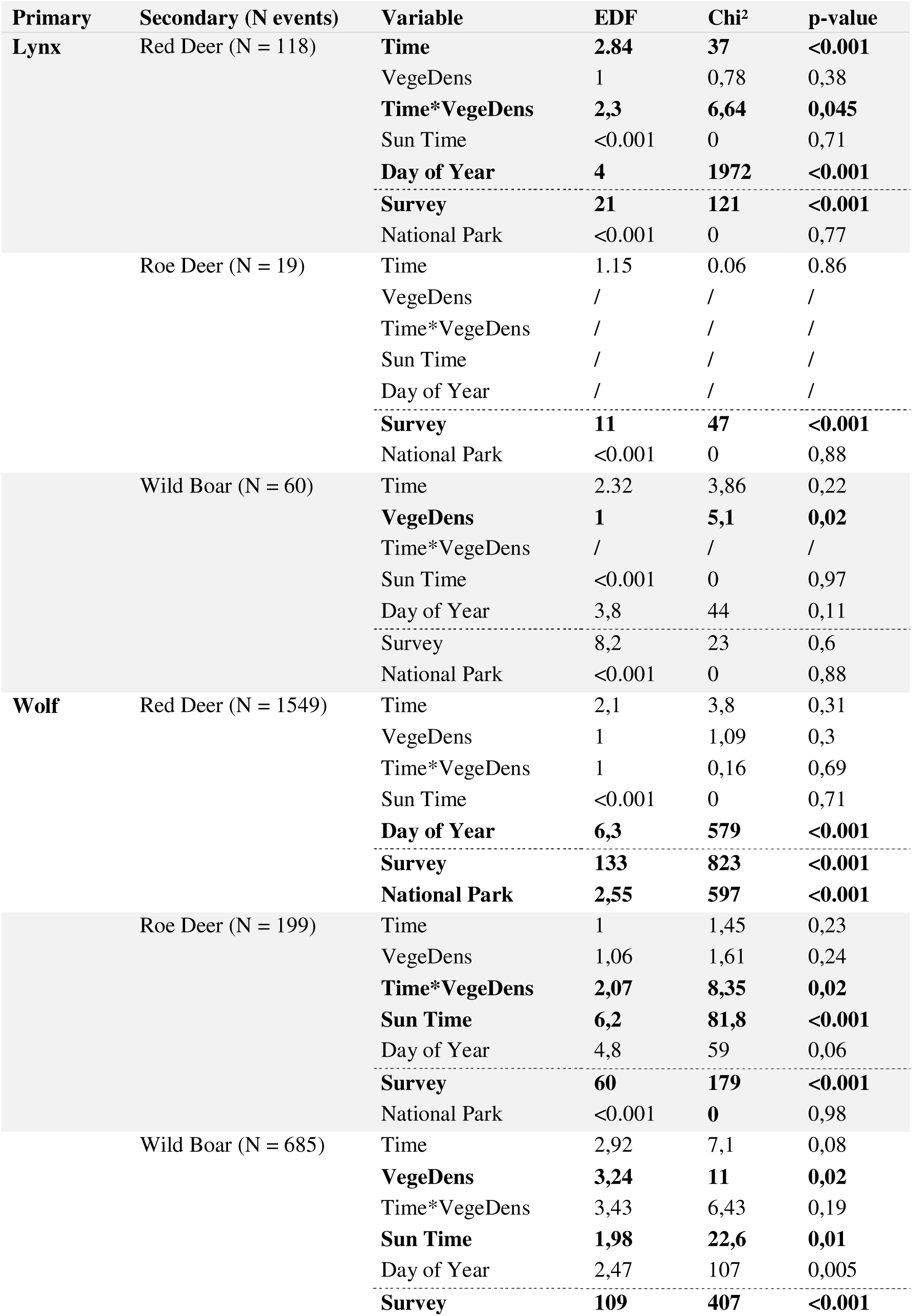
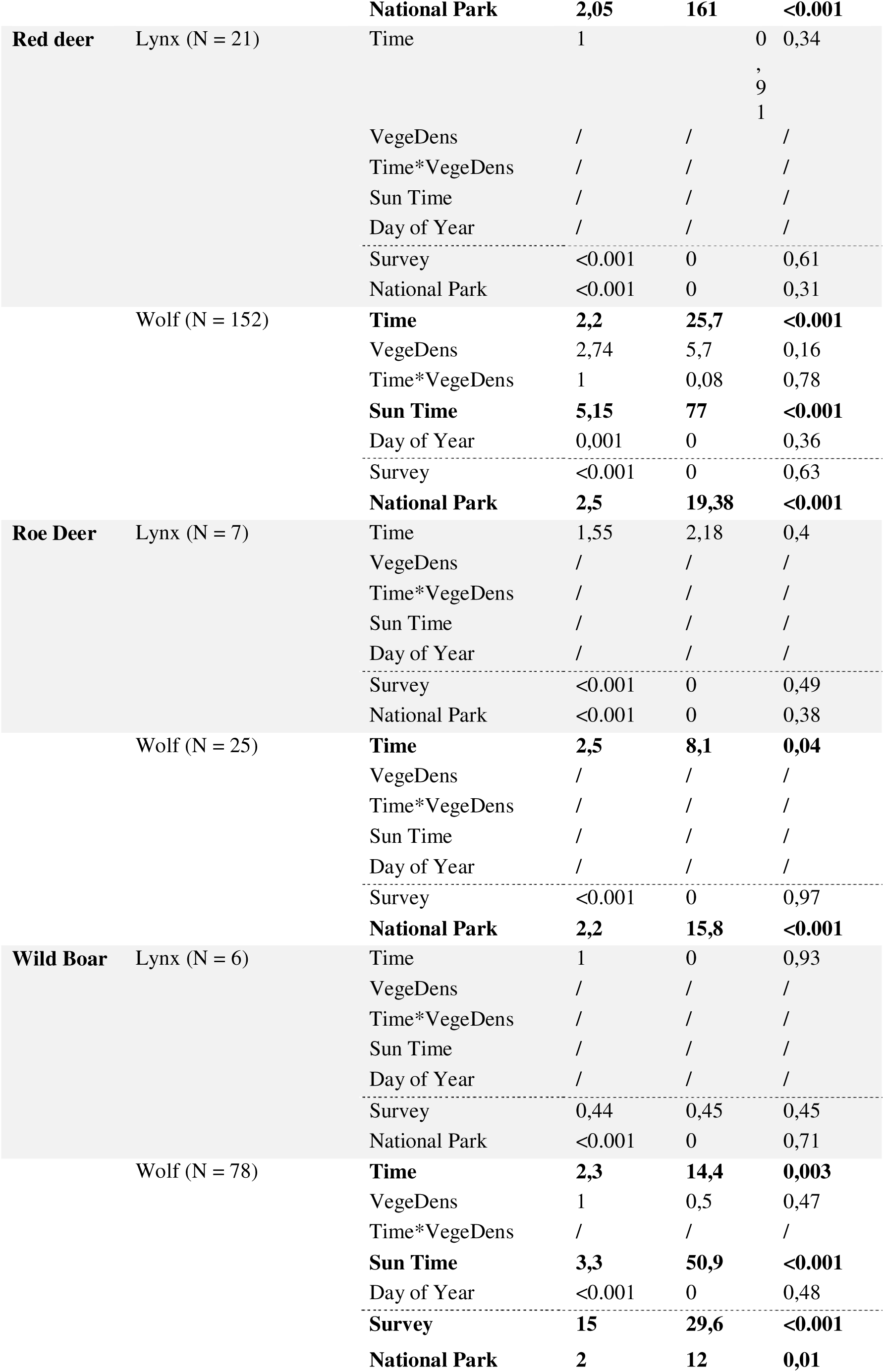
PAMM results for each primary-secondary combination include the effective degrees of freedom (EDF, it is not indicative of effect size or direction, e.g., positive). The associated p-value tests for non-linear effect of time, i.e. whether the smooth function significantly differs from a flat horizontal line. Time = time (in days) after the occurrence of the primary species, VegeDens = vegetation density, Sun Time = time of the day anchored around sunrise and sunset, Day of Year = day of year, Survey = random effect to include dependence between secondary recurrent event within the same survey after primary occurrence, National Park = random effect to include dependence of events within the same national park.Variables followed with the sign “/” were not included in the associated model due to sample size limitations.

## Results

Red deer were most frequently observed (n = 16276), followed by wild boar (n = 6980) and roe deer (n = 3832, Fig.4 bottom row displaying all four national parks number of post prey-survey). Survey numbers were primarily constrained by predator events (Fig. 4). Wolves were the most prevalent predators in the four protected areas (n = 269, comprising 152, 25 and 78 wolf recurrent events after red deer, roe deer and wild boar, respectively, Fig. 4), while lynx were only observed in two (n = 41, comprising 21, 7 and 6 lynx recurrent events after red deer, roe deer and wild boar, respectively, Fig. 2 & 4). This led to fewer post-lynx surveys and limited lynx recurrent events (Fig. 3 & 4). In contrast, there was ample information on post-wolf surveys and wolf recurrent events. The limited data on combinations involving roe deer, lynx, and wolves, or wild boar and lynx, prevented testing the effect of vegetation density for these prey-predator combinations (Table 3, Appendix 1).

**Figure 3:**
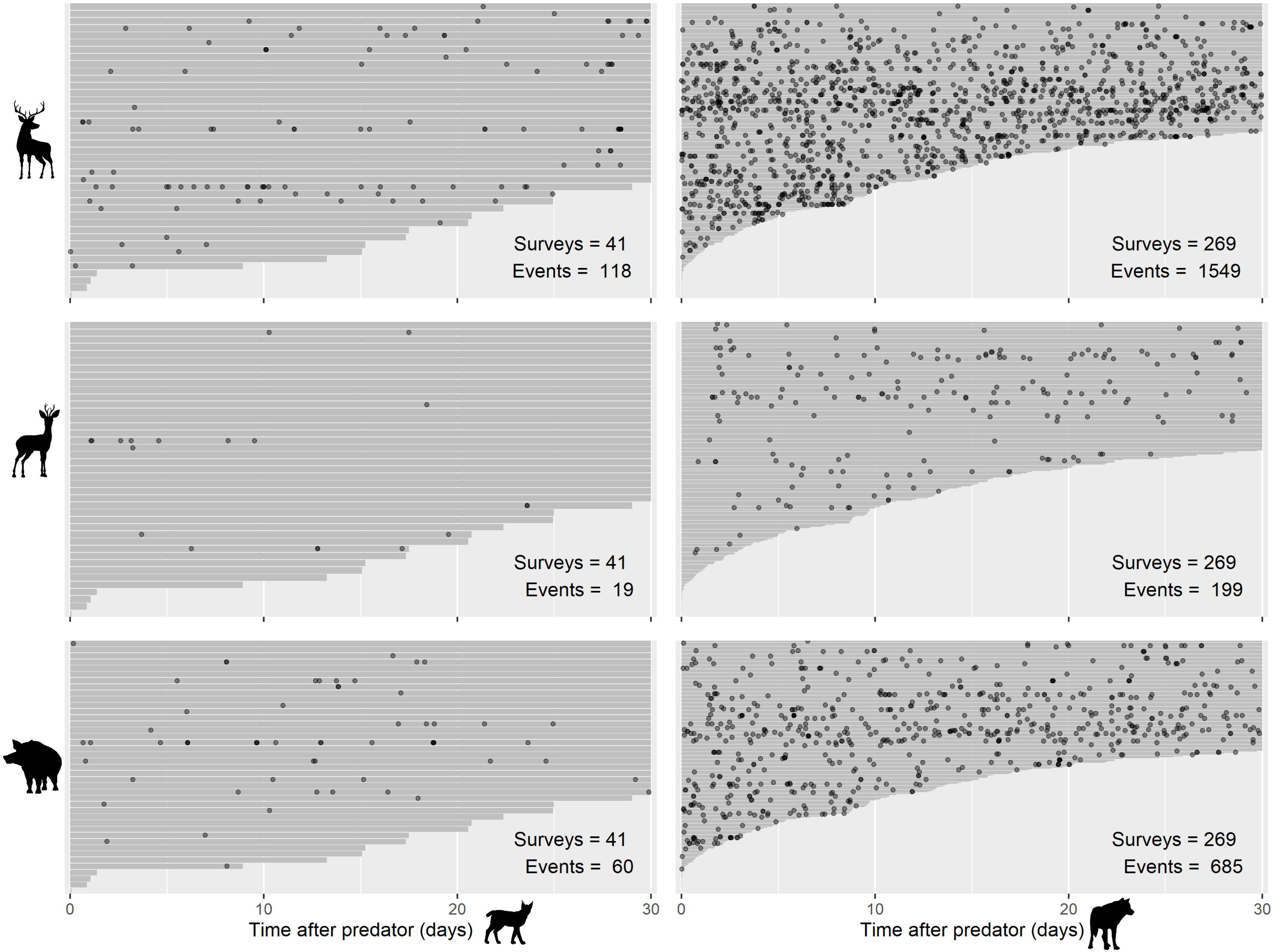
Graph of prey recurrent events within post-predator survey, structured by prey (rows) and predator species (columns, adapted from Chiou et al. 2021). Each grey line represents a survey, sorted by decreasing duration (max. 30 days). Black dots represent independent prey recurrent events within the surveys. The number of surveys and recurrent events per combination is shown for each plot. Rows, from top to bottom: red deer, roe deer, wild boar. Columns, from left to right: Eurasian lynx, wolf.

**Figure 4:**
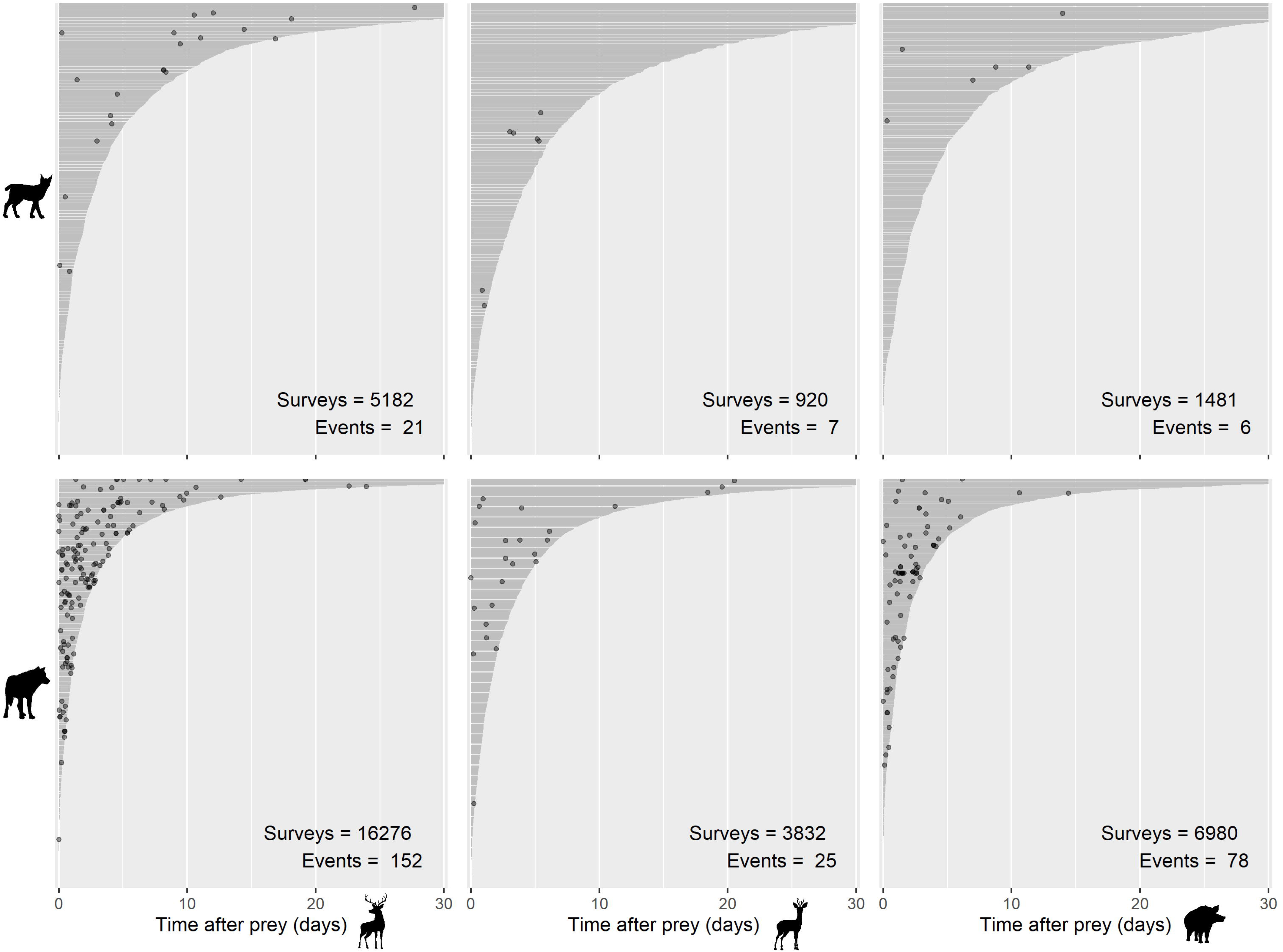
Graph of predator recurrent events within post-prey survey, structured by predator (columns) and prey species (rows, adapted from Chiou et al. 2021). Each grey line represents a survey, sorted by decreasing duration (max. 30 days). Black dots represent independent predator recurrent events within the surveys. The number of surveys and recurrent events per combination is shown for each plot. Rows, from top to bottom: Eurasian lynx, wolf. Columns, from left to right: red deer, roe deer, wild boar.

The prey response to predators showed notable changes only for red deer following lynx occurrences (Table 3). Red deer exhibited avoidance behaviour, with a slight increase in visitation rate after twenty days, which disappeared as vegetation density decreased (i.e. interaction effect), though variability was substantial (Fig. 5 & 7), and surveys after twenty days were limited (Fig. 3). Although the vegetation density was statistically significant regarding wild boar after lynx, and after wolf, as well as an interaction of time and vegetation density for roe deer after wolf (i.e., different from a horizontal flat line, Table 3), predictions were associated with high variability and no clear pattern was observed (Fig. 7). No patterns were observed for wild boar or roe deer (Table 3, Fig. 5), and no interaction effect with vegetation density was found for the combination wild boar following wolf (Table 3).

**Figure 5:**
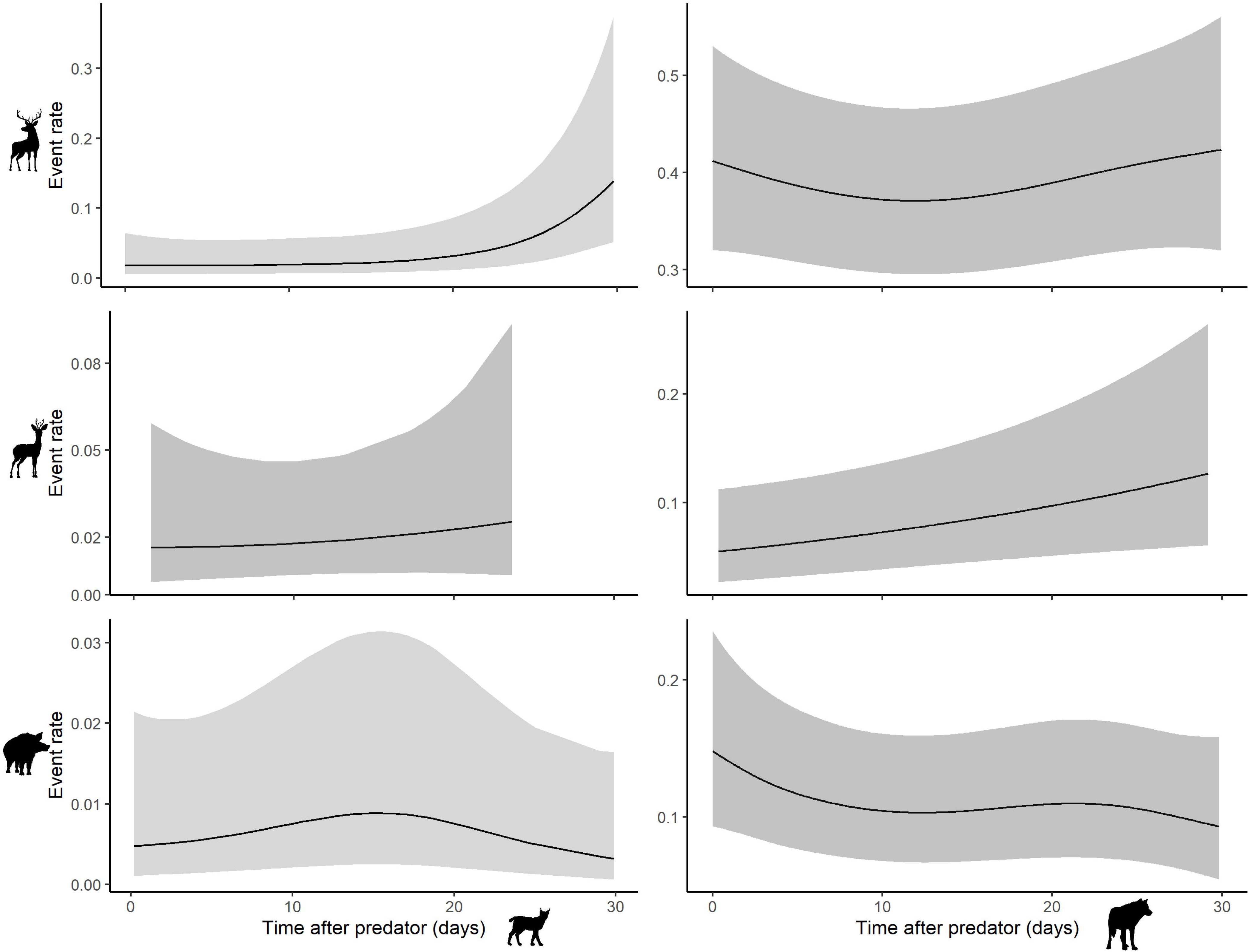
Estimated prey visitation rate over 30 days after predator occurrence. Columns represent prey species (red deer, roe deer, wild boar), and rows depict predators (Eurasian lynx, wolf).

Lynx showed no significant response to prey (Table 3, Fig. 6) likely due to the very low sample size (n = 41 in total, Fig. 4). Wolves exhibited visitation rates ca. 6.3, 4.6, and 5.2 times higher immediately after encountering red deer, roe deer, and wild boar, respectively, indicating strong initial attraction to all prey (Table 1, values approximated from differences in visitation rates between t = 0 day and either the plateau or the end of the smooth function estimation around t = 10 days, dotted horizontal lines Fig. 6). The variability in wolf visitation rate after fifteen days, following all three prey species, was due to limited surveys and few wolves’ recurrent events (Fig. 4 & 6). No wolf recurrent events occurred after fourteen days following wild boar encounters, precluding further trend estimation (Fig. 4 & 6). No effect of the vegetation density and its interaction with the time was observed (Table 3).

**Figure 6:**
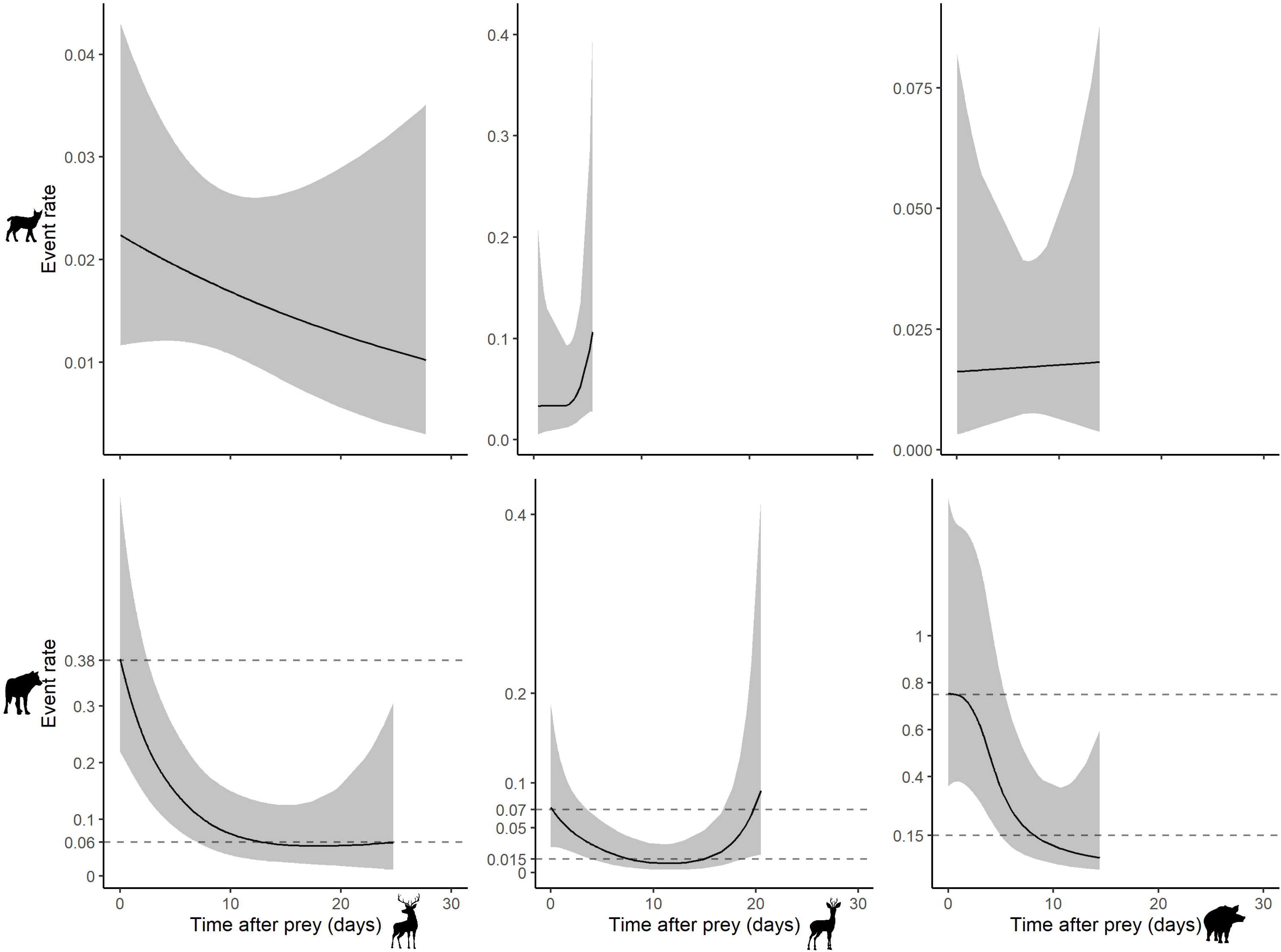
Estimated predator visitation rate over 30 days after prey occurrence. Columns represent predator species (Eurasian lynx, wolf); rows depict prey species (red deer, roe deer, wild boar). Horizontal dashed lines in the right column compare wolf visitation rates at t = 0 and t = 10 days. High variability after fifteen days in wolf estimates is due to limited surveys (Fig. 4).

**Figure 7:**
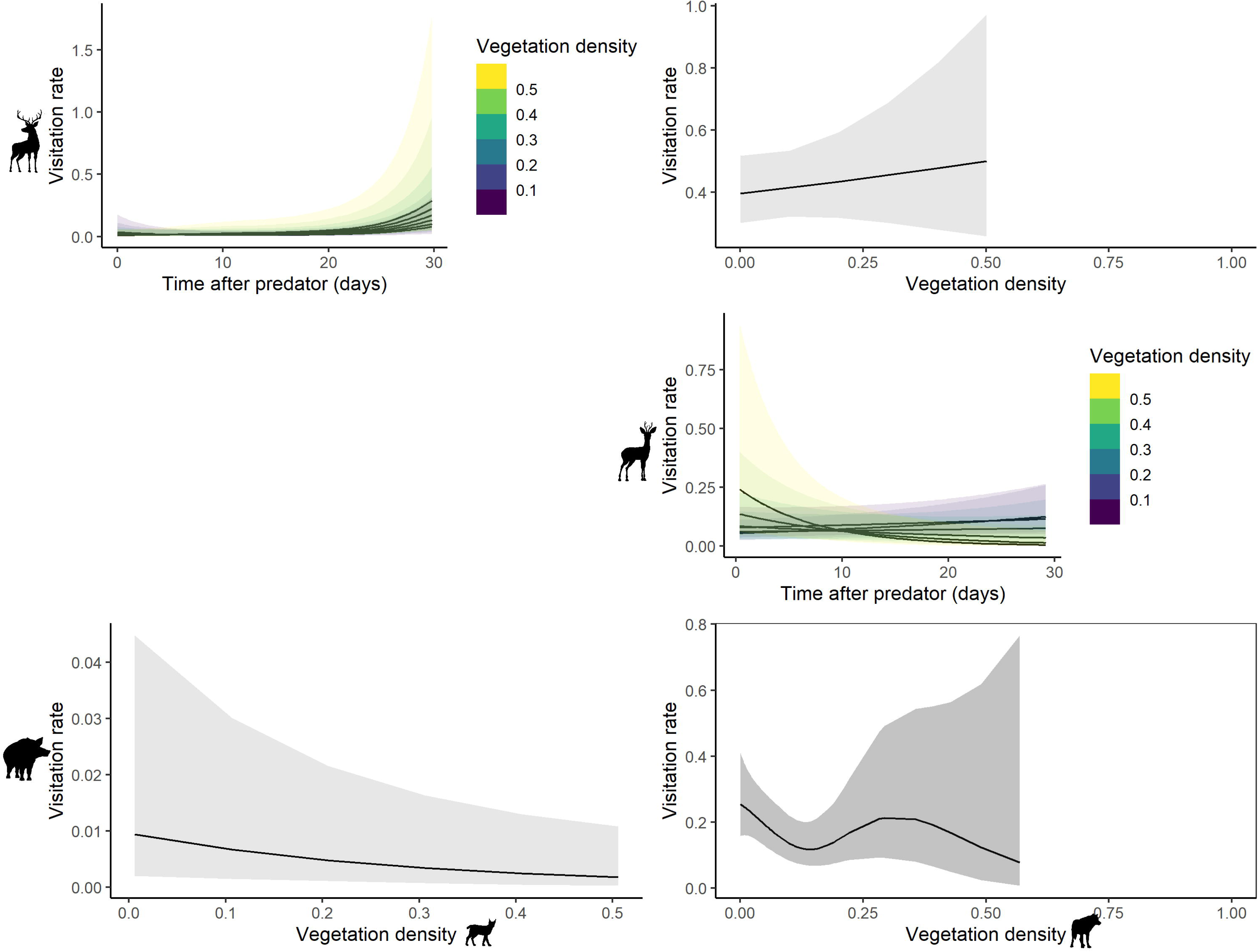
Estimated prey visitation rate either according to vegetation density alone (grey ribbons); or in interaction with the time after predator for the following combinations: red deer after lynx and roe deer after wolf. Columns represent predator species (Eurasian lynx, wolf); rows depict prey species (red deer, roe deer, wild boar).

**Figure 8:**
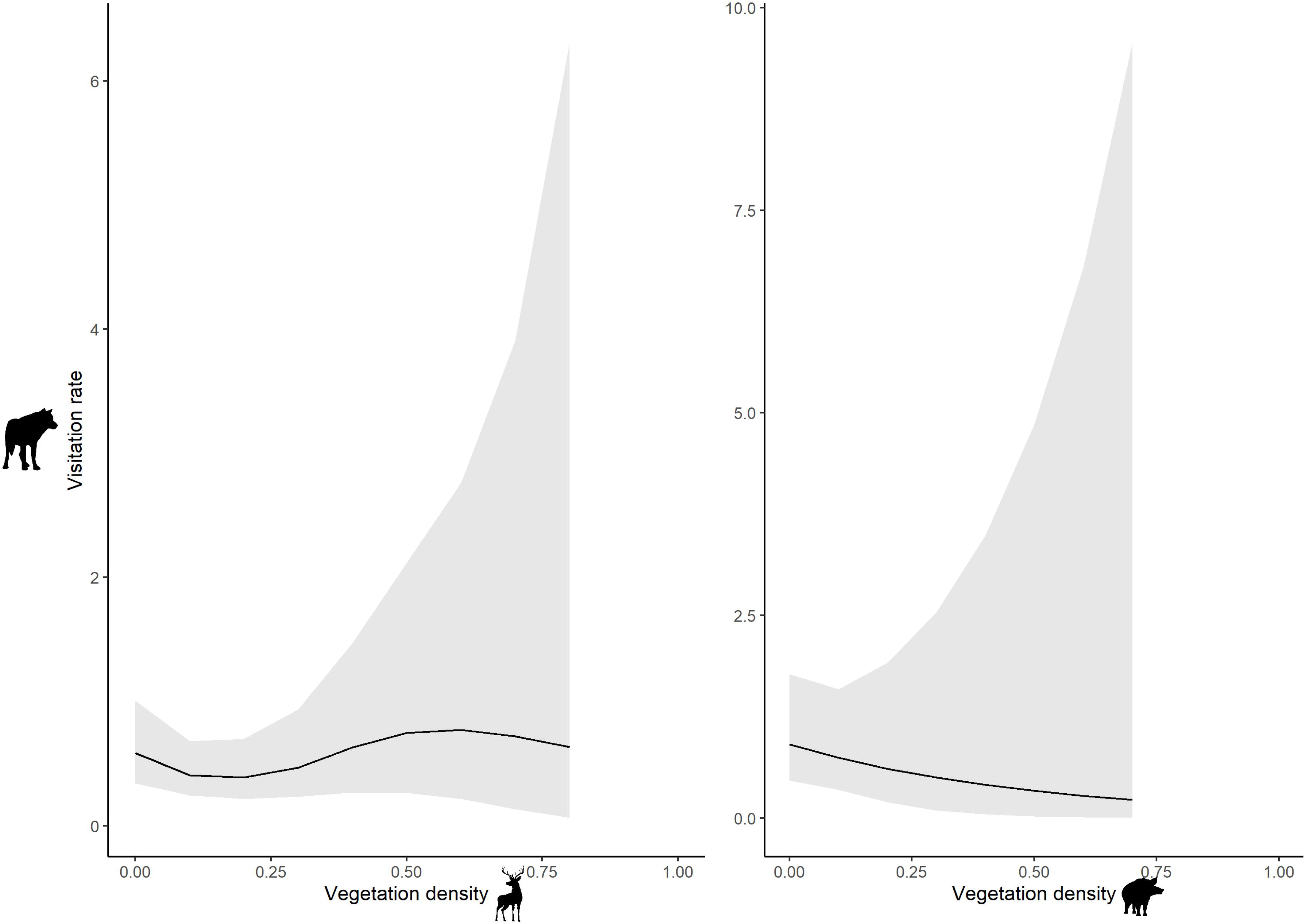
Estimated predator visitation rate according to vegetation density alone after red deer and wild boar occurrences (left and right column, respectively).

Results regarding the time of day (Sun Time) and the day of year (DoY) are presented in Table 3 and predictions in Appendix 2. We will not discuss the dynamics associated to these two control variables as i) they served as control variable and ii) were not purely representative of the behaviour of the secondary species but also of the primary species. For instance, there are peaks of events for red deer after wolf in May-June and September-October (Appendix 2), but this might be due to i) a true increase of red deer activity, ii) an increase of wolf activity leading to more post-wolf surveys at these periods or iii) a mix of both.

## Discussion

Our findings did not support the hypotheses of avoidance effects by grey wolves on all three prey species or by Eurasian lynx on roe deer and wild boar (H1, H2, H3). Only red deer showed a slight, enduring avoidance towards lynx. Conversely, our results supported the hypothesis that wolves respond to prey presence with a significant increase in visitation rates (H4). Lynx showed no clear response towards prey occurrence (H5). The limited number of lynx events after prey occurrences might be the reason for such an absence of the expected pattern. Placing camera traps on trails (Filla et al. 2017), where lynx is more active at night (Heurich et al. 2014), could improve sample size. Consequently, our discussion will focus on the wolves’ response. Finally, no clear pattern associated with the vegetation density was observed for prey (H6), and no effect was detected for predators (H7).

### Proximate co-occurrence on unmarked species

The study aimed to assess proximate co-occurrence, with an emphasis on understanding predator-prey interactions. However, studies using unmarked species and camera traps face a limitation: the absence identification at the individual-level. For example, when observing a flat visitation rate of red deer after wolf occurrences, it’s hard to distinguish between (i) the same individual red deer repeatedly visiting or (ii) several individuals appearing once and not returning. The first scenario suggests no response, while the second indicates avoidance. The reality is likely a mix of both, but distinguishing them is currently impossible. However, some signs can help infer the more likely scenario. Lower population density and smaller home ranges increase the likelihood of observing the same individual. For instance, given the red deer’s population density (2 indiv/km², Tourani et al., 2023), monthly home range (around 4.7 km²*, unpublished data*), and camera trap spacing (max. 1 trap/km²) in the Bavarian Forest National Park, observing dozens of different individuals at a site during a 30-day survey is unlikely, suggesting the first scenario of absence of response. Similarly, the low density of predators compared to prey, especially for gregarious species like wolves (e.g., Mech & Peterson, 2003), makes it more likely to observe the same individual or pack.

### The predation sequence

While the focus was not on providing a complete approximation of encounter rates, by considering also spatial and temporal overlaps, some interesting conclusions can nonetheless be drawn. Wolves showed expected attraction patterns towards prey, supporting our hypothesis H4. However, prey did not show the anticipated avoidance patterns, as expected under the landscape of fear theory. We suggest three reasons behind this asymmetry in reactive response.

First, predators and prey face different pressures throughout the predation sequence. Among all components (i.e. spatio-temporal proximity, encounter, ignorance or avoidance post-encounter, capture or escape from attack), prey can optimize specific components of their response to avoid predation, without necessarily addressing all the other components, even completely ignoring some of them. Some may specialize in avoiding risky habitats, while others rely on early detection through vigilance, grouping, or alternative defensive mechanisms (Gaynor et al., 2019, Gerber et al., 2024). For instance, van Beeck Calkoen et al. (2021) did not observe change of visitation frequency and vigilance duration in presence of wolf’s urine but highlighted reduced visitation duration. Predators, while certainly not always maximizing their predation rate (e.g. when resting, drinking, territory exploration, etc.), must however achieve a minimum success probability in each component when attempting to complete the predation sequence. Thus, prey responses may seem counterintuitive (e.g., Hebblewhite et al., 2005; Kauffman et al., 2007; Cuzack et al. 2017; Prat-Guitart et al., 2020; Sunde et al., 2022) but would reflect this flexibility of managing predation risks, showcasing the complexity of predator-prey interactions.

Second, this absence of response from prey might be related to the value of the information on the current location of a predator. In their spatially explicit individual-based model mimicking predator-prey space use and foraging, Fraker & Luttberg (2012) observed that as movement and perception of predator increased relatively to those of prey, the latter being more neutral toward predator location. Given that a predator could be anywhere, the information of where they were the step before had no value and prey were then selecting for high-quality patches rather than avoiding predators. In our study, given that wolf are large-territory animals (95% estimated AKDE Home Range = □50; 570] km², Vorel et al., 2024) compared to their prey, and able to move fast, the cues of previous wolf presence might not be as informative as they would from lynx, for instance.

Finally, in Ferry et al. (2024), the PAMM as well as the others tested approaches could not efficiently detect (i.e., risk β < 0.2) weak or short-term effect (e.g. <12 hours) despite hundreds of recurrent events available. Therefore, we could not completely rule out the avoidance of prey towards predators, particularly regarding the response of roe deer towards wolf, but this avoidance might be too short or too weak to be detected solely based on camera trap data and the current statistical approaches.

### The effect of the environment and human

Dormann et al. (2018) stressed the importance of considering underlying processes and confounding factors when interpreting species interactions from co-occurrence data. Multi-Species Occupancy Models (MacKenzie et al. 2004) and joint Species Distribution Models (Pollock et al. 2014) can help by accounting for covariate effects on species occurrence. In spatio-temporal analyses, issues like a carcass causing apparent attraction between scavengers or herbivores gathering at waterholes during the dry season (Ferry et al. 2016) can lead to misinterpretations. Time-to-event analyses often struggle to incorporate covariates, but Prat-Guitart et al. (2020) used a survival framework as an exception. PAMM, given enough data, allowed us to examine vegetation density as a proxy for perceived predation risk and its impact on species interaction. Our analysis found no influence of this parameter on interactions. This may be due to the limited scope of our measurement, confined to a 4m radius around camera traps, covering about 64 m² whereas animal decision-making might encompass habitat information at a smaller scale. Additionally, camera trap placement often avoids dense vegetation to maximize detection probability, which might not accurately reflect the broader habitat context, hindering precise assessment of vegetation density effects. Finally, the random placement of camera traps in the forest limited our ability to capture human activities such as non-lethal recreation, which takes mostly place on trails, and hunting. Both lethal and non-lethal activities can impact the spatio-temporal activity of both predator and prey as well as their interactions (Peters et al., 2025; Courbin et al., 2022; Bonnot et al., 2020; Gaynor et al., 2018). Often, detailed data on human activities is scarce, limiting studies and methodology to address this topic. However, due to the marked and widely recognised impact human activities have on wildlife behaviour, incorporating human presence in data collection and analysis would be beneficial to understand predator-prey interactions in Europe’s human dominated landscape more holistically.

### A limited top-down control from large predators?

Our results show that when predator and prey visit the same site, prey do not exhibit short-term avoidance detectable at the spatio-temporal resolution of camera traps. This limited behavioural response implies that the local reduction in browsing pressure may be weaker than expected if one infers predator effects solely from visit dynamics. Such a pattern aligns with recent work showing that other behavioural dimensions, such as reduced visit duration, can mediate risk responses even when visitation rates remain unchanged (van Beeck Calkoen et al., 2021). In recolonising European landscapes, where wolves and lynx are often expected to restore top-down control on herbivores, these findings highlight that predator return does not automatically translate into strong suppression of site use by prey, and that managers should account for the multifaceted nature of fear-driven responses when anticipating impacts on browsing pressure.

## Conclusion

Our study delved into the spatio-temporal dynamics of interactions among three herbivore species (red deer, roe deer, and wild boar) and two carnivores (wolf and lynx) using camera trap data collected across four protected areas. Leveraging PAMM, we precisely estimated the temporal patterns of visitation rates for each species following the occurrence of another. Contrary to our initial expectations, none of the herbivores exhibited discernible reactions to the presence of predators. Conversely, we observed a notable attraction of wolves to all three prey species. This stark contrast suggests an inherent asymmetry in predator-prey interactions: while predators must navigate through all stages of the predation sequence to successfully achieve a predation event, prey species have the flexibility to evade predation by succeeding either in all or only a portion of these stages. These intriguing findings highlight the potential existence of asymmetry in predator-prey interactions, warranting further investigation.

## Acknowledgements

The present publication is a partial result of the R&D project ‘Ungulate monitoring in German National Parks’ (FKZ: 3518 83 0200), which was supported by the Federal Agency for Nature Conservation (BfN). We deeply thank Volker Spiecher from Müritz National Park and Jürgen Stein from Sachsenforst State Company, Nature Reserve Administration Königsbrücker Heide / Gohrischheide Zeithain for providing data.

## Statement of authorship

NF conceived this idea; NF led the writing of the manuscript; NF led analyses; CF, AP & AB carried out the investigations, CF and MH acquired funding and conceived the data collection design. All authors contributed critically to the drafts and gave final approval for publication.

## Data accessibility statement

R scripts and data are available from the Zenodo Digital Repository https://doi.org/10.5281/zenodo.13911909.

