## Appendix 2 for "Prey attracting but not avoiding predators suggests an asymmetric investment in the predation sequence"

### Appendix 2 – Sun Time and Day of Year results

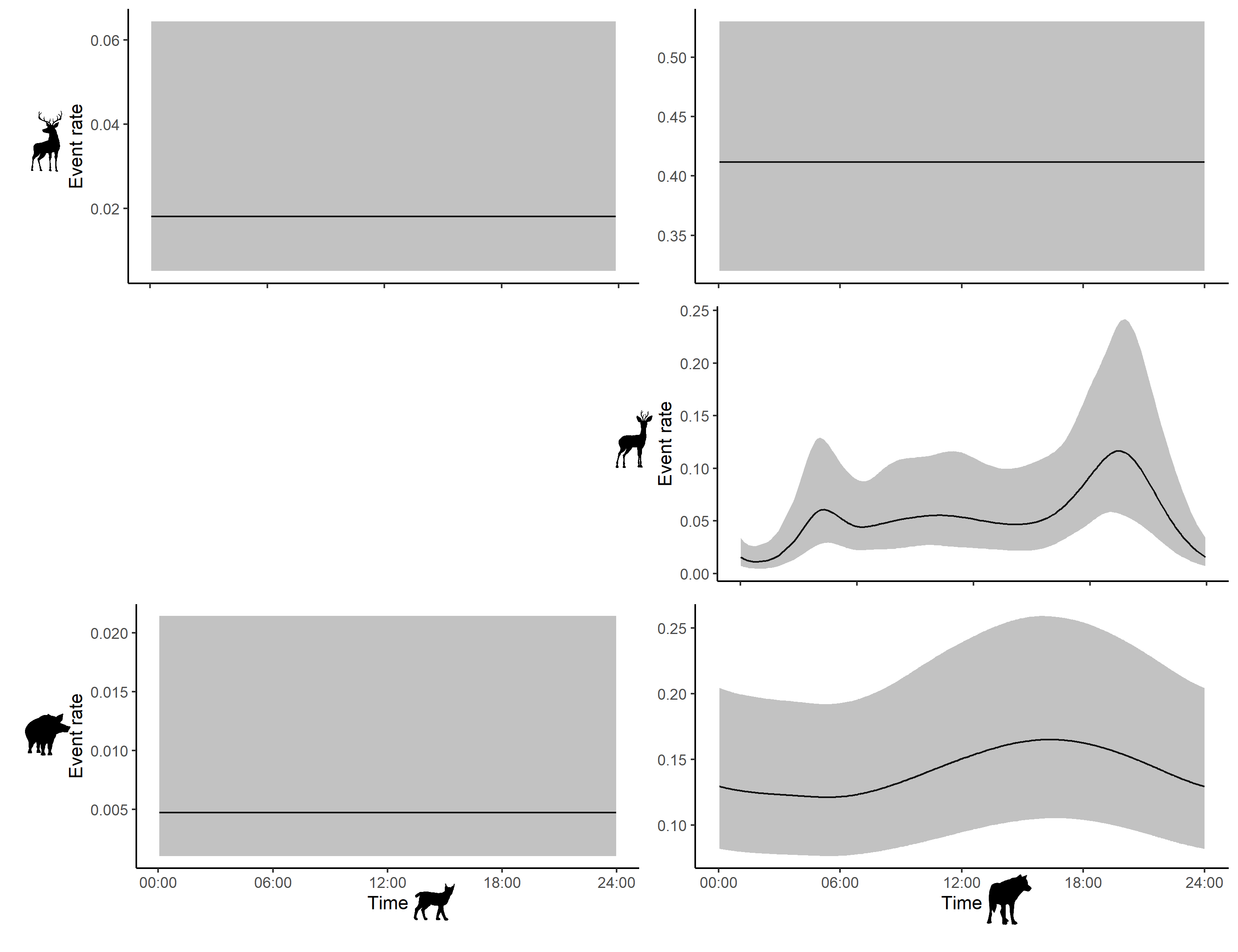

Figure S2-1: Estimated prey visitation rate over the time of the day (with average anchoring transformation). Rows represent prey species (red deer, roe deer, wild boar) as secondary species, and columns depict predators (Eurasian lynx, wolf) as primary species.

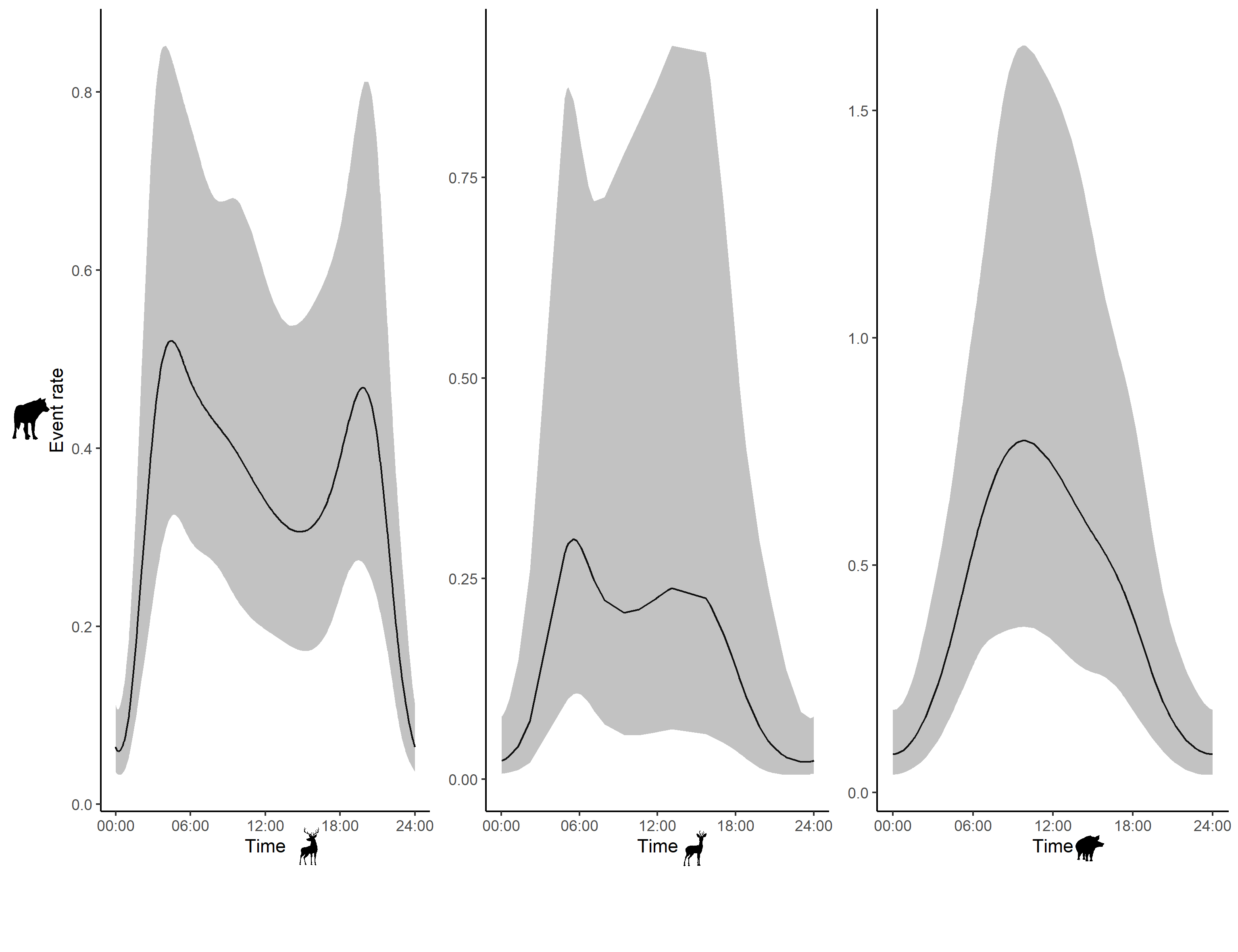

Figure S2-2: Estimated predator visitation rate over the time of the day (with average anchoring transformation). Row depict predator (wolf) as secondary species, and columns represent prey species (red deer, roe deer, wild boar) as primary species.

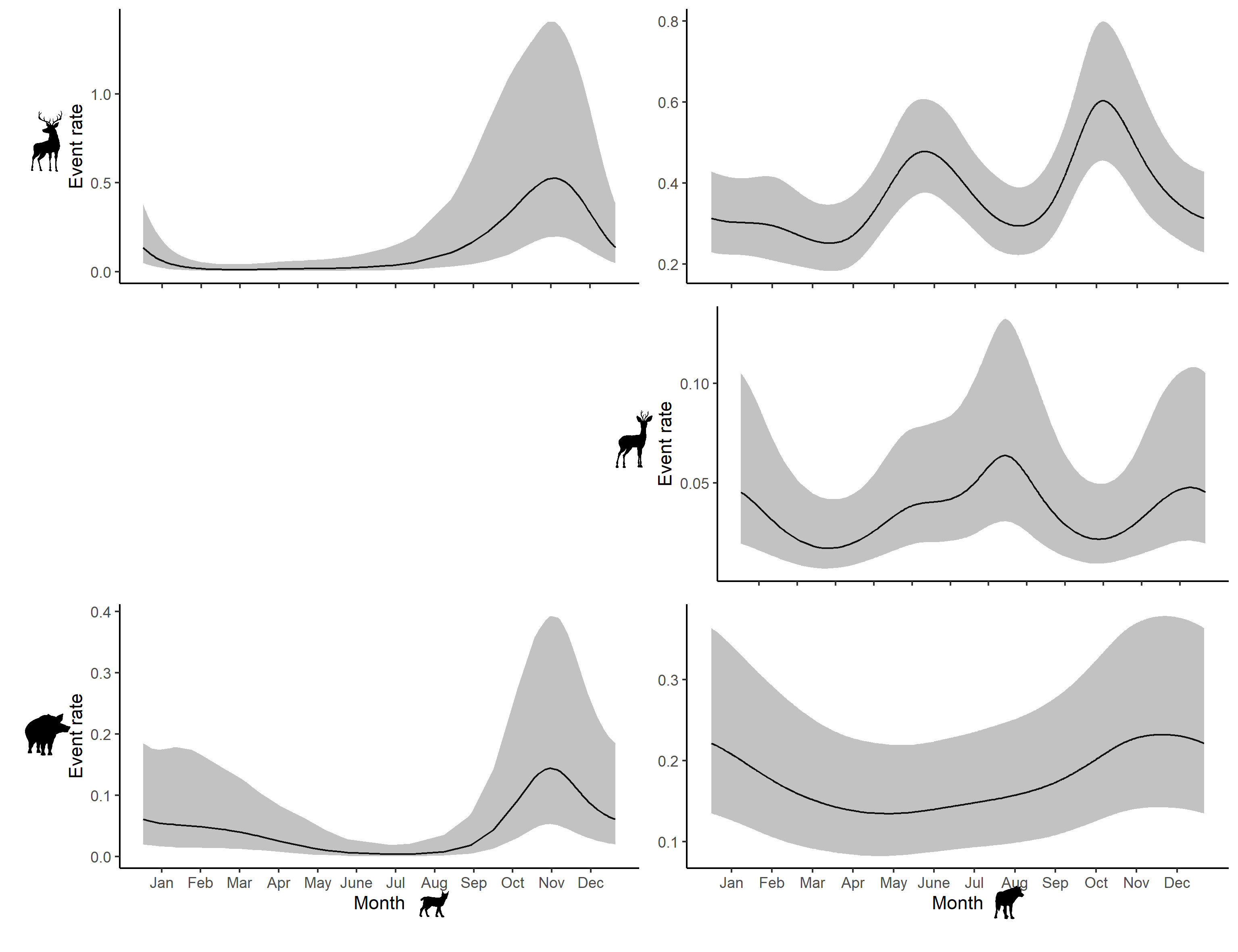

Figure S2-3: Estimated prey visitation rate over the day of the year. Rows represent prey species (red deer, roe deer, wild boar) as secondary species, and columns depict predators (Eurasian lynx, wolf) as primary species.

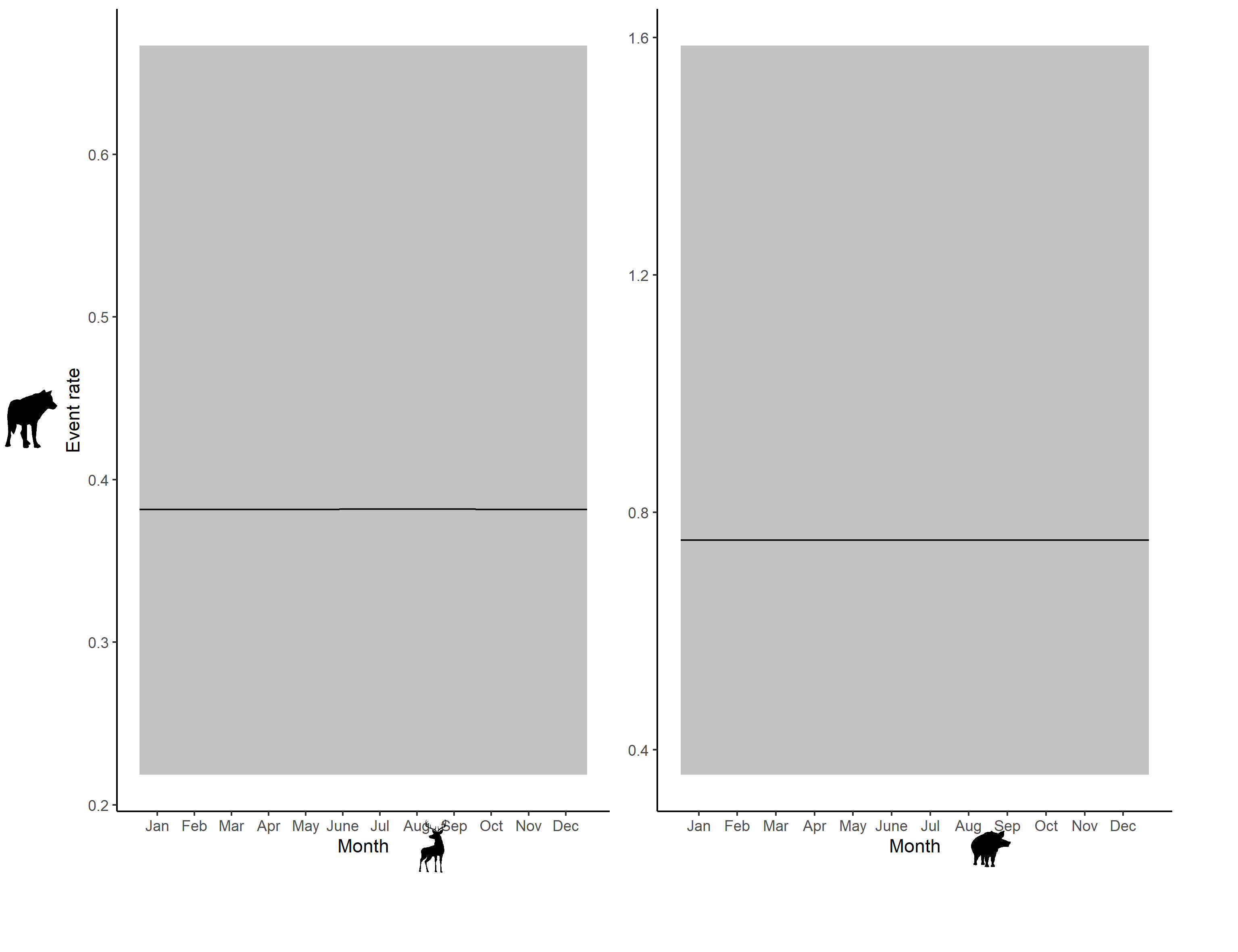

Figure S2-4: Estimated predator visitation rate over the day of the year. Row depict predator (wolf) as secondary species, and columns represent prey species (red deer, roe deer, wild boar) as primary species
