## Appendix 1 for "Prey attracting but not avoiding predators suggests an asymmetric investment in the predation sequence"

### Appendix 1 – Survey definition and censoring

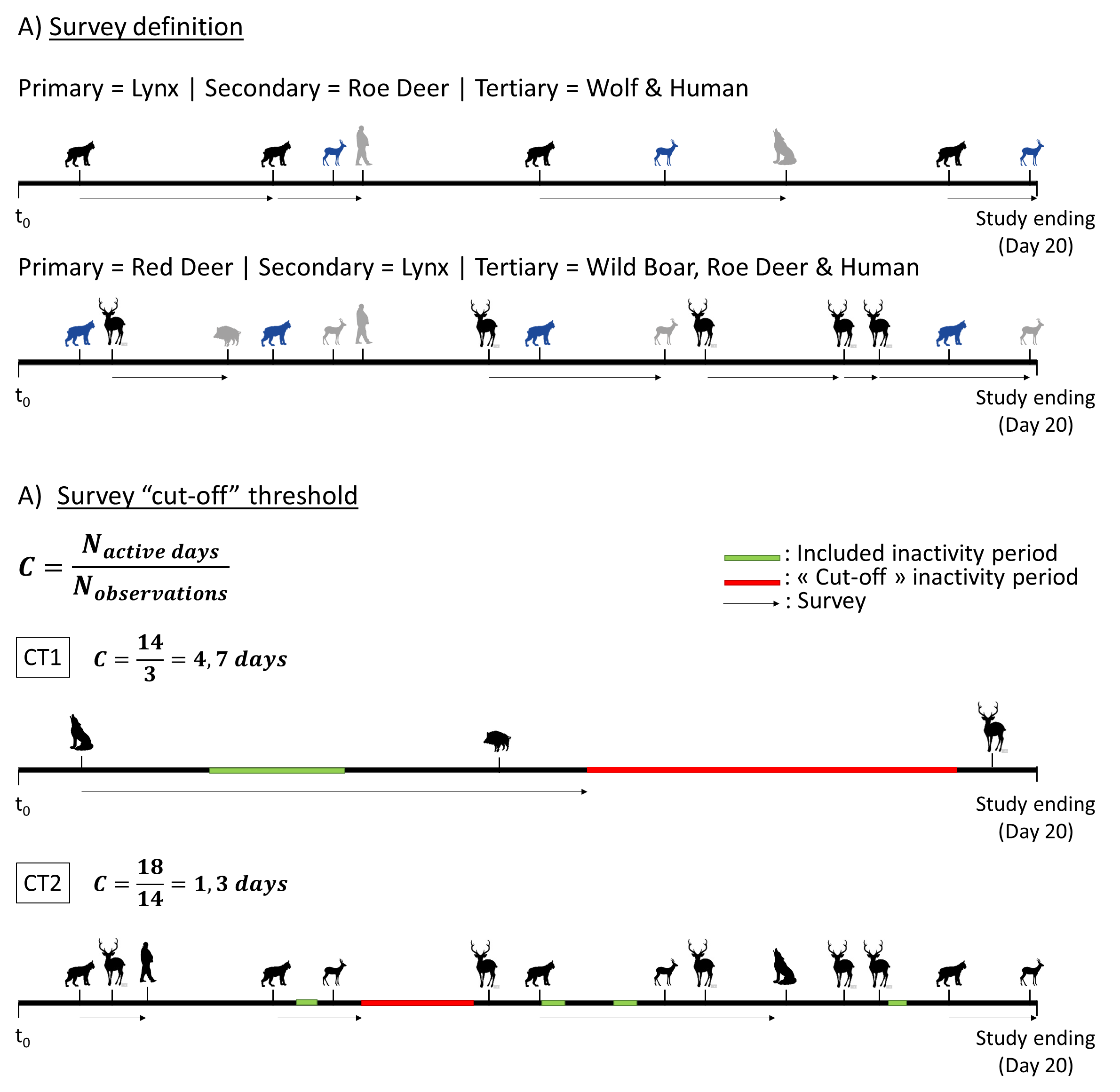

**Figure S1**: A) Graphical representation of survey definition and censoring. Horizontal lines depict hypothetical camera trap monitoring at two sites (CT1 and CT2) over 20 days each. Silhouettes represent species events. A) Survey construction based on the primary (black silhouette), secondary (blue silhouette) and tertiary species (grey silhouette). A survey is initiated by a primary occurrence at a camera trap site and is denoted by a black arrow. Surveys conclude due to i) a censoring species observation (primary or tertiary), ii) reaching the survey duration threshold, iii) the study's end, or iv) a cut-off inactivity period (see after). B) Cut-off inactivity periods were defined for each camera trap site as any duration surpassing the average days between two animal events (e.g., 4.7 days and 1.3 days for CT1 and CT2, respectively).
